# FveOCRs - a database for open chromatin prediction in wild strawberry based on a large language model

**DOI:** 10.1101/2025.10.22.683913

**Authors:** Muzi Li, Yuzhu Wang, Zhennan Zhao, Stephen M. Mount, Qing Ma, Zhongchi Liu

## Abstract

**Background:** Woodland strawberry (*Fragaria* vesca) is a widely used model system for cultivated strawberries and Rosaceae for molecular genetic studies. Nevertheless, available databases for its cis-regulatory element identification are limited. With the emergence of large language models in plant research, strawberry research could benefit significantly from applying these models. One such model derived from Plant DNA Large Language Models (PDLLMs) effectively predicts open chromatin regions (OCRs), which are nucleosome-depleted areas typically associated with cis-regulatory elements. However, improvement of the model’s accessibility and utility will be needed to facilitate the model’s application in strawberry as well as other plant species.

**Results:** We developed the FveOCRs database (http://liulab.online:33838/FveOCRLiuLabTestVer25), which predicts open chromatin regions located within 5,000 bp upstream sequences, the first and second introns and exons, and the longest introns in *Fragaria vesca* genes. Predictions were generated using a sliding-window approach based on one of the PDLLMs. In the database, users could predict open chromatin regions for input DNA sequences from any plant species and obtain visual graphs with precise predicted regions of open chromatins.

**Conclusions:** The FveOCRs database is the first database for open chromatin region prediction in *Fragaria vesca* based on a Plant DNA Large Language Model. It can also predict OCRs in user-provided DNA sequences from any plant species. The resource and its accessible visual graphs will facilitate discovery of cis-regulatory elements and engineering of gene expression in strawberry and other plant species.

## Background

Strawberry is one of the most economically important fruit crops worldwide. Global strawberry production has increased from 0.75 million tonnes in 1961 to 10.485 million tonnes in 2023 (https://www.fao.org/faostat/en/#home). Its popularity among consumers is driven by its attractive color, fragrant aroma, and diverse nutrients, including vitamins and antioxidants (1). The woodland strawberry (*Fragaria vesca*), a diploid species with a high-quality genome assembly and annotation, serves as a model system for the cultivated octoploid strawberry (2). It’s widely used to investigate the genetic mechanisms underlying key biological processes and traits and to facilitate genome editing for precision breeding. One key research area in *F. vesca* as well as other crops is to dissect regulatory mechanisms of different biological processes and metabolic pathways. Hence, the ability to predict, isolate, and manipulate cis-regulatory elements is at the forefront of plant synthetic biology and will benefit greatly from new tools, especially AI-based tools.

Open chromatin regions are nucleosome-depleted areas that can be accessed by trans-acting factors (3). These regions dynamically change according to environmental and developmental conditions (3) and typically contain cis-regulatory elements, including promoters, enhancers, and silencers, that recruit regulatory proteins to control transcription (4). Several AI-based computational tools have been developed to predict open chromatin regions in plants. For example, CharPlant (Chromatin Accessible Regions for Plant, https://github.com/Yin-Shen/CharPlant) (3) is a convolutional neural network-based model trained on datasets from four plant species, while PDLLMs (Plant Large Language Models, https://github.com/zhangtaolab/plant_DNA_LLMs) (5) comprises a suite of models designed for multiple tasks, including open chromatin prediction.

With the rapid advancement of sequencing technologies and computational methods, an increasing number of databases have been developed for *Fragaria vesca*, providing valuable genomic resources for this valuable model. The Genome Database for Rosaceae (GDR, https://www.rosaceae.org/) (6) is a well-maintained and widely used repository that hosts multiple versions of genomes and annotations for Rosaceae species and provides a range of tools and viewers. The Strawberry eFP Browser (https://bar.utoronto.ca/efp_strawberry/cgi-bin/efpWeb.cgi) (7) enables users to visualize gene expression across diverse *Fragaria vesca* tissues and developmental stages. The *Fragaria vesca* Co-expression Network Explorer (http://liulab.online:33838/fvesca/) (8) provides robust consensus co-expression networks allowing researchers to identify co-expressed genes that may be functionally related. The Rosaceae Fruit Transcriptome Database (ROFT, https://www.rosaceaefruits.com/) (9) supports comparative analysis of ortholog expression across four Rosaceae species in various tissues during early fruit development. In addition, the Plant Chromatin Accessibility Database (PlantCADB, https://bioinfor.nefu.edu.cn/PlantCADB/) (10) identifies accessible chromatin regions using experimental data from high-throughput chromatin accessibility assays in 37 plant species, including *Fragaria vesca*. However, the *Fragaria vesca* datasets in PlantCADB are limited to a few tissues and developmental stages, and use the older version genome FraVesHawaii_1.0 released in 2010.

Despite these genomic resources, no AI-based database currently exists for open chromatin prediction in *Fragaria vesca*. Here, we developed the *Fragaria vesca* Open Chromatin Regions (FveOCRs) database, a user-friendly platform that integrates a sliding-window approach with the plant-dnamamba-BPE-open_chromatin model derived from PDLLMs (5). By stepwise scanning of DNA sequences in defined windows, this approach enhances the precision of OCR prediction and enables users to explore the results through graphs and tables. We also experimentally validated a positive cis-element within an upstream OCR for *FveMYB10*. Identifying open chromatin regions will facilitate the discovery of cis-regulatory elements, thereby enabling the precise control of gene expression through genetic engineering of these elements.

## Construction and content

### Database development

In *Fragaria vesca*, the 5,000 upstream regions of genes, the first and second exons and introns, and the longest introns were scanned using a sliding-window approach (step size: 10 bp; window sizes: 200, 500, and 1,000bp). Open chromatin prediction was based on the *Fragaria vesca* Genome 4.0.a2 (11). The 5,000 upstream regions corresponded to the sequences located 5,000 bp upstream of the gene transcription start sites (TSSs) specified in the gene feature rows of the GFF annotation file. The exon and intron sequences were derived from the isoforms encoding the longest proteins. For each sliding window, the fine-tuned plant-dnamamba-BPE-open_chromatin model from PDLLMs (5) was applied to predict the probability of the region being a full, partial or non-open chromatin region. The plant-dnamamba-BPE-open_chromatin model is built on Mamba (12), a new large language model architecture based on State Space Models (SMMs), and employs byte-pair encoding (BPE) (13) as its tokenizer.

The prediction results were deposited in MongoDB (v5.0.31). The database was developed with the R package Shiny (v1.10.0) (14), and communication between MongoDB and the Shiny application was supported by the R package mongolite (v4.0.0) (15). The home page graph was generated by BioGDP (https://biogdp.com/) (16). The genome browser embedded in the database was generated using the R package JBrowseR (v0.10.2) (17). The database was deployed on an Ubuntu operating system (v24.04).

### Database structure and components

The FveOCRs database comprises four main functional tabs, “OCR Prediction (Fve)”, “JBrowse”, “Retrieve Data”, and “OCR Prediction (Any Plant Seq)”, and three supplementary tabs “Home”, “Help”, and “Others”) (Figure 1). The details of these tabs are described below.

**Figure 1.**
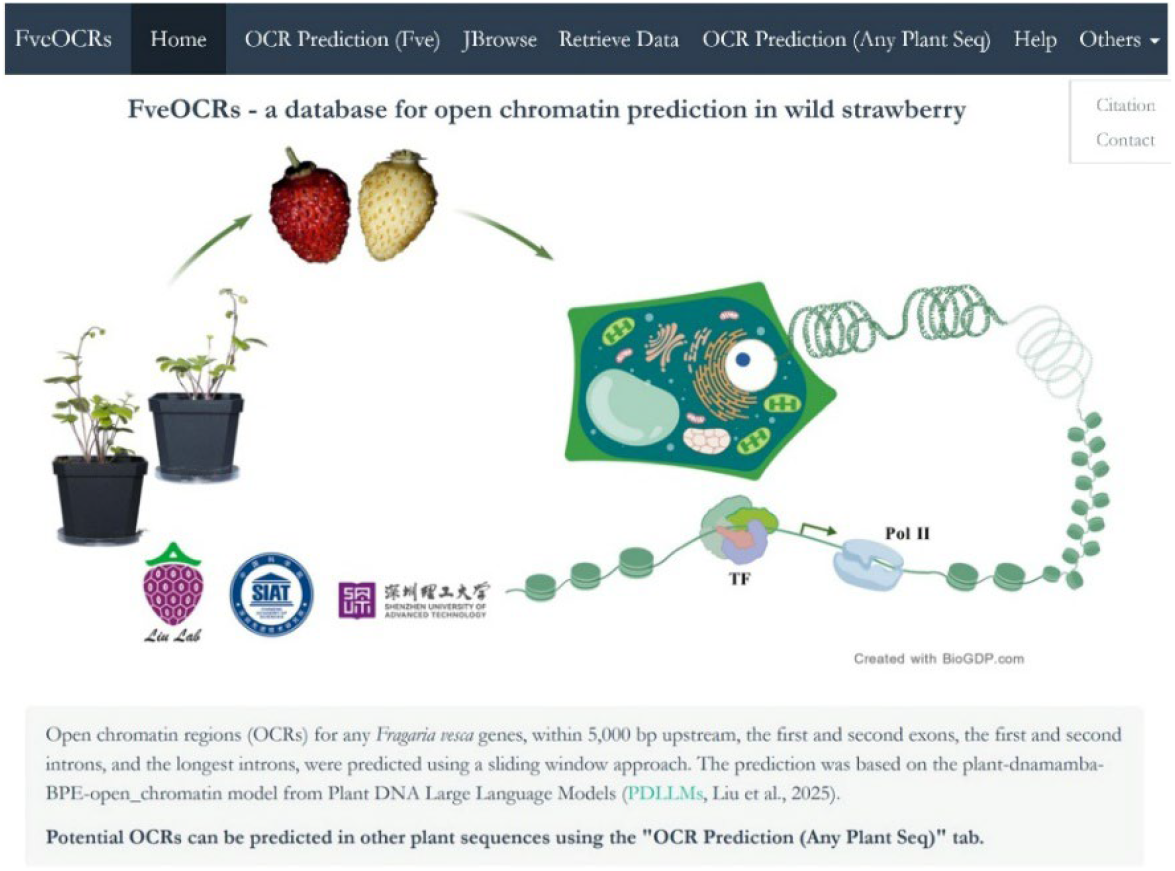
The “Home” tab of the FveOCRs database. In addition to “Home”, there are six tabs on the top panel, “OCR Prediction (Fve)”, “JBrowse”, “Retrieve Data”, “OCR Prediction (Any Plant Seq)”, “Help”, and “Others”.

The “**OCR Prediction (Fve)**” tab (Figure 2) allows users to search for open chromatin regions in the gene of interest. In the search box (Figure 2A), users can enter a gene ID (*Fragaria vesca* Genome v4.0.a2 (11)) in the Gene ID box and select the query type (5,000 bp upstream, the first and second exons, the first and second introns, or the longest intron). These sequences were chosen because they often contain cis-regulatory elements. One then chooses the sliding window size (200 bp, 500 bp, or 1,000 bp). The sliding window moves from 5’ to 3’ in the gene’s direction by 10 bp at each step (Figure 2B). The prediction of OCR for each window was displayed in the scatter plots, showing the probability of each window being a fully open chromatin, a partially open chromatin, or a non-open chromatin (Figure 2C). Each dot in the scatter plots represents one window. An explanation is provided alongside the plots to aid the interpretation. To achieve fast speed, we pre-computed the OCR probabilities with the plant-dnamamba-BPE-open_chromatin model from PDLLMs (5) and stored the results in MongoDB. Additionally, beneath these plots is information on the gene’s chromosomal location, the coordinates of the selected region, and the best Arabidopsis BLAST hit.

**Figure 2.**
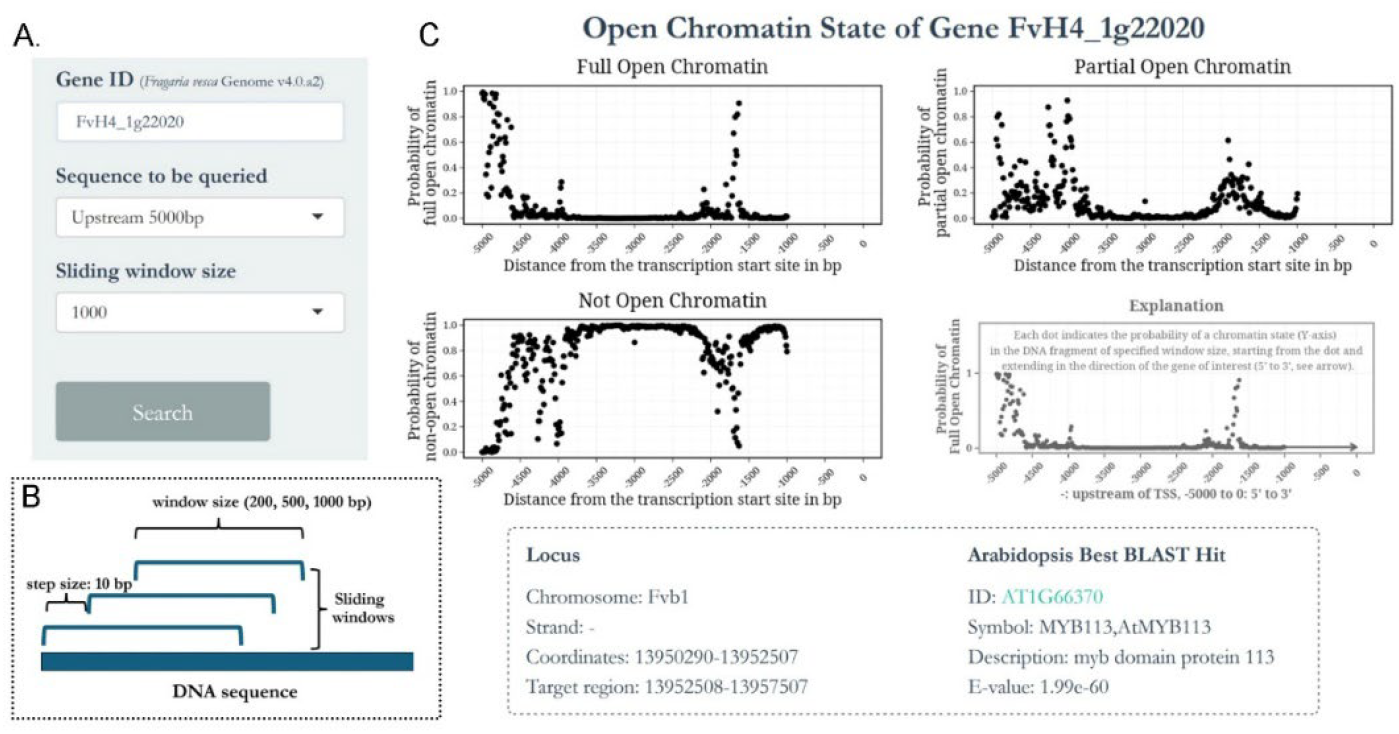
Illustration of the “OCR Prediction (Fve)” tab. (A) Search box used to define query gene ID, corresponding gene’s sequence for analysis, and the window size. (B) Illustration of the sliding window approach (not displayed in webpage). (C) The three scatter plots (top left, top right, and bottom left) respectively display the probabilities of sliding windows being full, partial, or non-open chromatin regions. The fourth plot (bottom right) uses full open chromatin prediction data as an example to demonstrate how to interpret the points and axes in the scatter plots. The locations of the gene and target region in the *F. vesca* genome v4.0.a2, as well as the gene’s best Arabidopsis Blast hit, are shown beneath the scatter plots.

The “**JBrowse**” tab (Figure 3) provides an alternative visualization approach, where the genome browser for *Fragaria vesca* Genome v4.0.a2 (11) is integrated with the OCR prediction outcome. Users can select multiple prediction tracks to view the probabilities of sliding windows at full, partial or non-open chromatin state (Figure 3A). One important note is that the OCR prediction probabilities, shown as blue lines, in one track are for genes on the same DNA strand. Users can choose both + and - tracks to visualize OCR predictions for genes on both strands. For example, the tracks “U5000_s10W500_+_FullOCR” and “U5000_s10W500_-_FullOCR” (Figure 3B) provide OCR probability for genes on the positive strand and negative strand, respectively.

**Figure 3.**
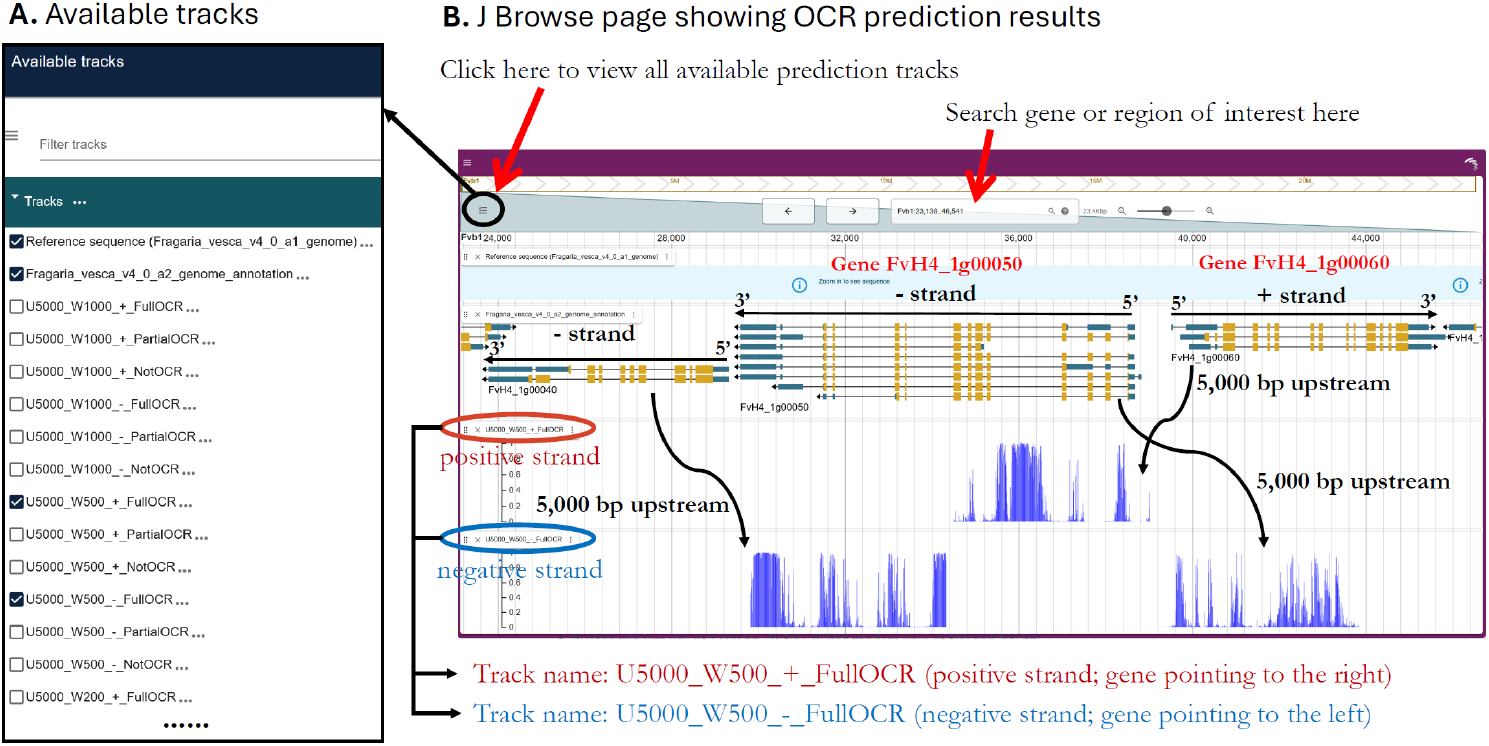
Illustration of the “JBrowse” tab and interpretation of the results. (A) Different tracks available for the users. It is visible when one clicks the upper left conner “three short lines” symbol (as shown by the red arrow). (B) The genome browser display of OCR prediction. Users can search for a gene or genomic region using the box at the top of the browser (red arrow). Each blue line indicates the OCR probability for a window, starting at the blue line and extending in the gene’s (5’ to 3’) direction. Both positive and negative strand tracks are shown. Two adjacent vertical lines are 10 bp apart. The *F. vesca* genome v4.0.a1 with annotation v4.0.a2 was used.

Similar to the “OCR Prediction (Fve)” tab, the chosen DNA sequence was scanned with a window size of 200bp, 500bp or 1000bp and at a 10 bp shift toward the gene’s 3’ end (Figure 2B), yielding probability scores (blue lines) for every shift. The “JBrowse” tab allows users to examine nearby genes, which is particularly useful when open chromatin regions overlap with adjacent genes. For instance, a predicted OCR upstream of the gene FvH4_1g00060 (positive strand) is located within the intron of the nearby gene FvH4_1g00050 (negative strand) (Figure 3B), suggesting that the intron region may serve dual functions - alternative splicing on the negative strand and transcriptional regulation on the positive strand.

The “**Retrieve Data**” tab (Figure 4) enables users to enter a gene ID, choose a region and a window size, and retrieve a table with pre-calculated probabilities indicating whether sliding windows in the selected region are classified as full, partial or non-open chromatin regions. Users can download the table and corresponding sequences with the download button. This tab allows users to obtain the precise DNA sequence within each window.

**Figure 4.**
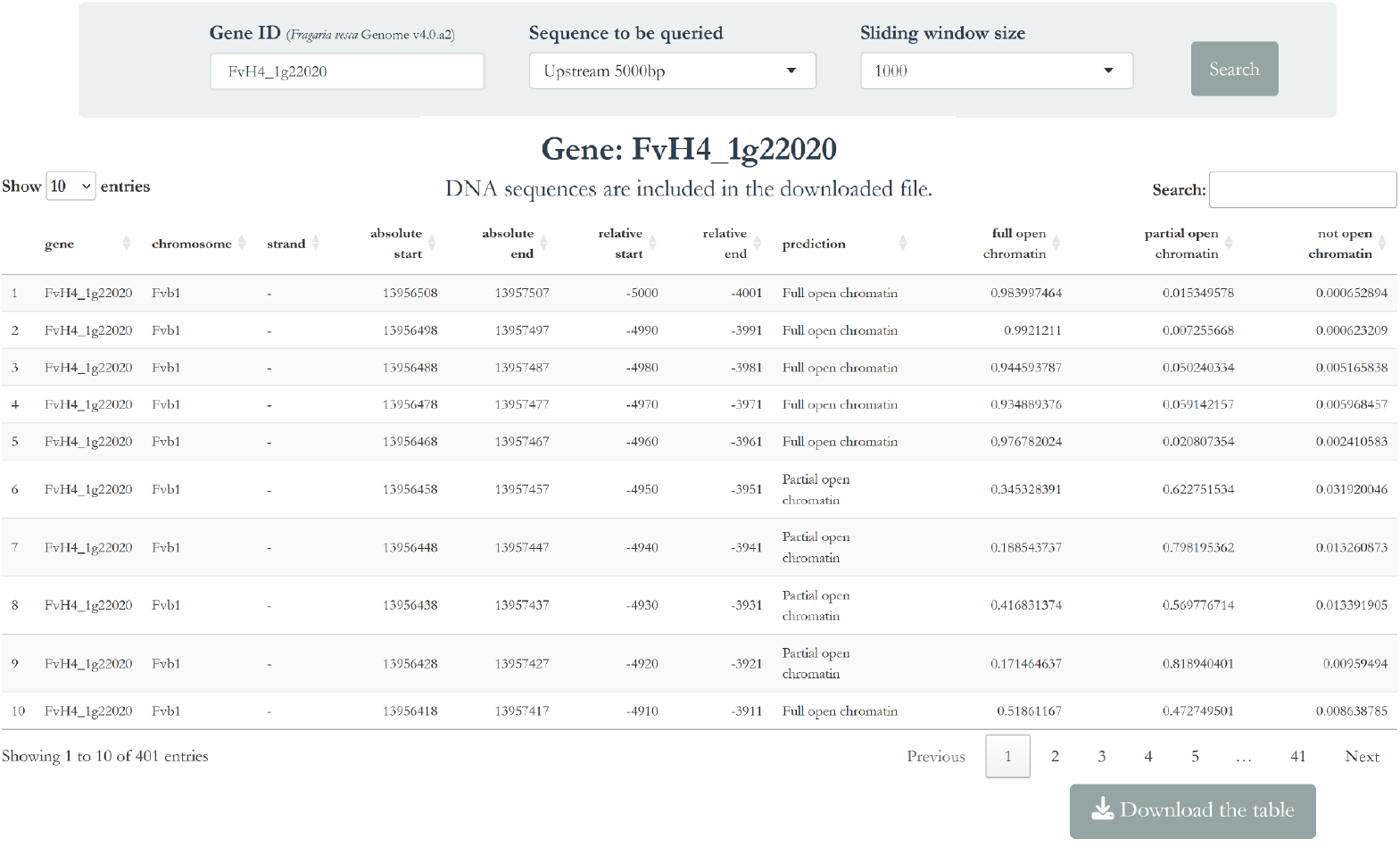
Illustration of the “Retrieve Data” tab. The table provides users with data on the predicted probability of each sliding window (each row) being a full, partial, or non-open chromatin region. It also includes the chromosomal location of each window and its relative position to the transcription start site (TSS), exons or introns, depending on the selected region. The downloaded table further contains the DNA sequence corresponding to each sliding window.

The “**OCR Prediction (Any Plant Seq)**” tab (Figure 5) expanded the utility of this database beyond wild strawberries. It allows users to predict open chromatin regions of any input plant sequences. Specifically, users input a piece of DNA sequence that can come from any plant species and click the “Predict OCRs” button, which will generate the scatter plots in real time (Figure 5). The prediction is similarly based on the plant-dnamamba-BPE-open_chromatin model from PDLLMs (5) with the sliding-window approach. Input sequence is limited to a length up to 2kb (<=2,000 bp), which will be scanned with the user-defined step and window size. The “Download the prediction” button provides the probability table along with the window sequences.

**Figure 5.**
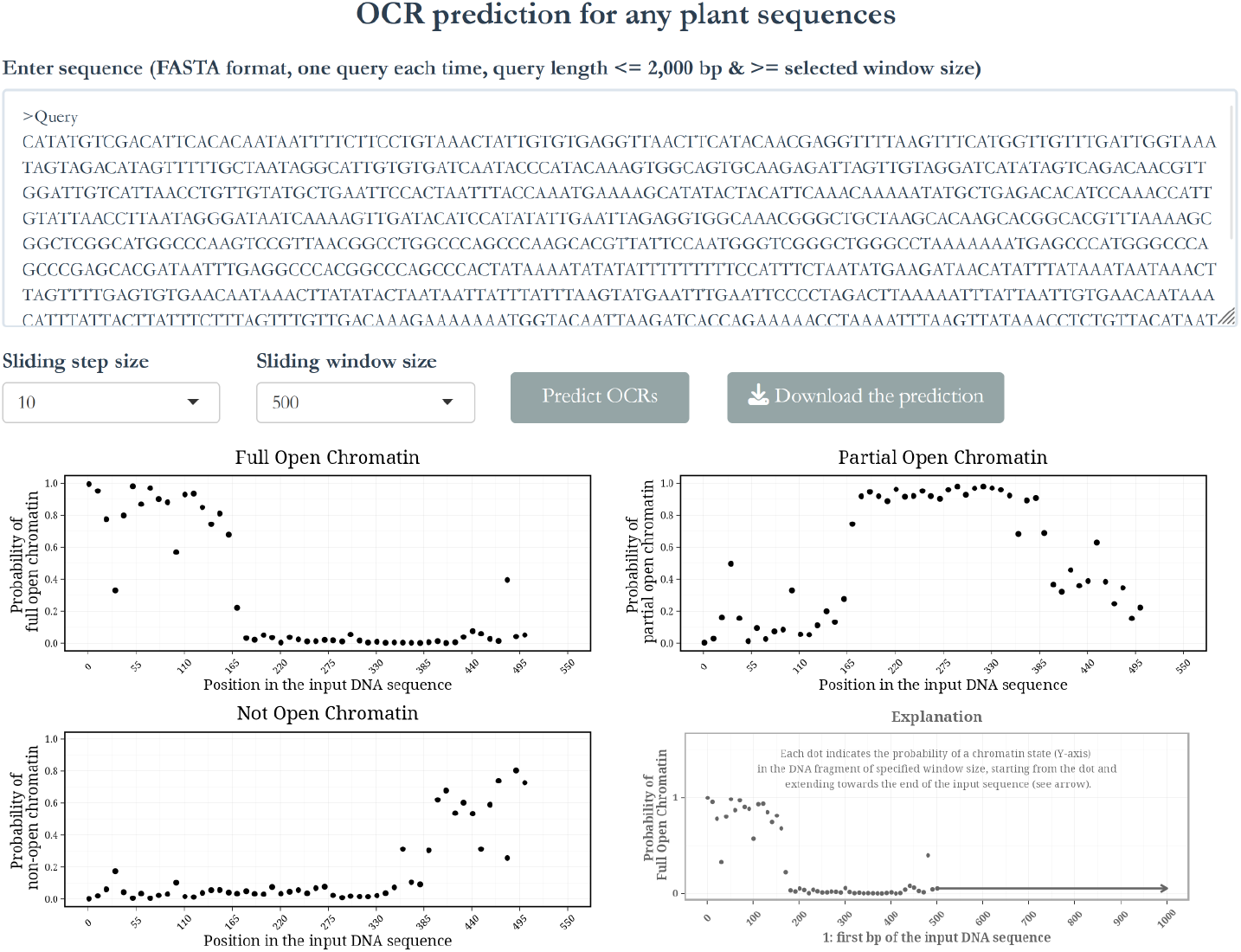
Illustration of the “OCR Prediction (Any Plant Seq)” tab. A single-stranded DNA sequence of up to 2.0 kb can be entered into the query box. Users can then select the sliding step size and window size from the pull-down menu beneath the query box and click “Predict OCRs”. The results will be presented as scatter plots, which may take a moment to appear since they are generated in real time.

In addition to the functional tabs, three supplementary tabs provide further details about the database. The “**Home**” tab presents an overview of the database. The “**Help**” tab offers comprehensive guidelines and illustrations to aid users in using the database and interpreting the outputs. The “**Others**” tab provides instructions for citing the database and the email address and GitHub information for contacts.

## Utility and discussion

### Model evaluation

The emerging AI-models in biological studies are reshaping biological research and accelerating research progress. The recently published Plant Large Language Models (PDLLMs) (5) offered a variety of fine-tuned models for predicting functional genomic elements in plants, including open chromatin regions, core promoters, and histone modification regions. Among the PDLLMs is a model “plant-dnamamba-BPE-open_chromatin”, which was trained on ATAC-seq, DNase-seq, Faire-seq, and MNase-seq datasets from diverse plant species, tissues and developmental stages, including *Fragaria vesca*. Hence, plant-dnamamba-BPE-open_chromatin is an important model for open chromatin prediction in plants.

To assess the accuracy of the plant-dnamamba-BPE-open_chromatin model (5) in predicting OCRs, we tested this model on experimentally determined MNase hypersensitive sites (MHSs) that correlate with open chromatin sites. These 96,841 MHSs identified in *Fragaria ananassa* ‘Royal Royce’ (18) have a length ranging from 50 bp to 1,703 bp (Figure 6A). As they were not included in the pre-training or fine-tuning of the plant-dnamamba-BPE-open_chromatin model, this evaluation provides an independent validation of its prediction power. When we applied the model to test these 96,841 MHS sites, 63.716% of the MHS sites were predicted as full open chromatin, and 19.731% were predicted as partial open chromatin (Figure 6B), culminating in 83.447% of these MHSs as full or partial OCRs. Hence, the test demonstrated that the plant-dnamamba-BPE-open_chromatin model successfully predicted the majority of the OCRs even in a plant species outside its training set.

**Figure 6.**
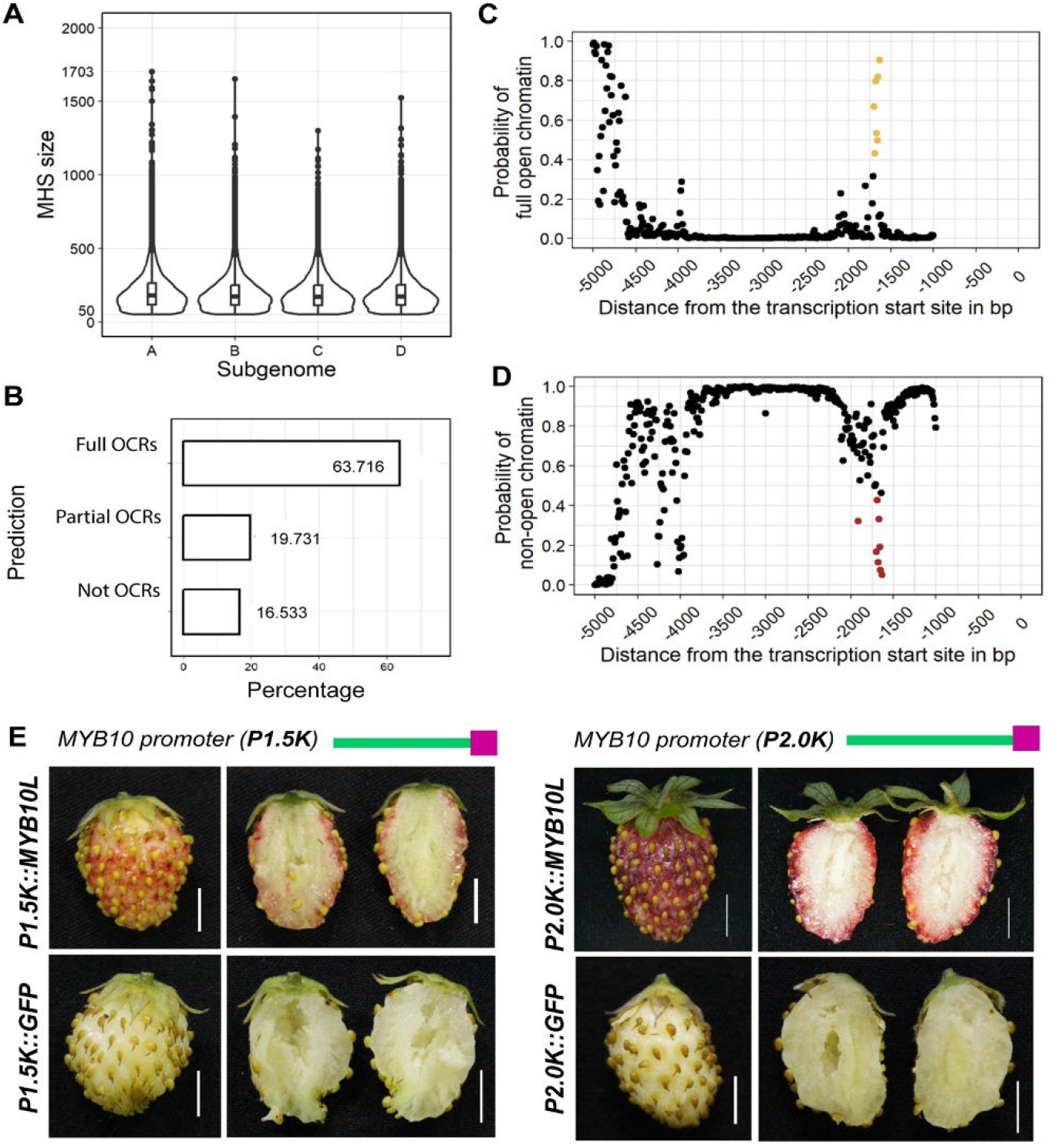
Validation of the plant-dnamamba-BPE-open_chromatin model and a case study with *FveMYB10* upstream sequence. (A) Summary of a prior experimental data on the MNase hypersensitive sites (MHSs) across the subgenomes (A, B, C, D) of a cultivated strawberry (*Fragaria ananassa* cv. Royal Royce). (B) Percentage of cultivated strawberry MHSs classified as full, partial, or non-open chromatin regions by the plant-dnamamba-BPE-open_chromatin model. (C) FveOCRs prediction of “Full Open Chromatin” in the upstream 5,000 bp region of *FveMYB10* (step size: 10 bp, window size: 1000 bp). Yellow dots indicate full open chromatin regions. (D) FveOCRs prediction of “Non-Open Chromatin” in the upstream 5,000 bp region of *FveMYB10* (step size: 10 bp, window size: 1000 bp). Brown dots full or partial open chromatin regions. (E) Transient overexpression of reporter gene *FveMYB10L* in *Fragaria vesca* fruit driven by the 1.5 kb (left) or 2.0 kb (right) upstream sequences of *FveMYB10*. GFP is used as a non-pigment control reporter. Scale bars: 0.5 cm.

### Platform improvement

In the prior publication, in addition to the PDLLMs command-line tools, a user-friendly online platform was made to facilitates functional DNA prediction (https://finetune.plantllm.org/) (5). However, the input sequence length and context, even shifted by a few bp, could dramatically alter the prediction outcome by the plant-dnamamba-BPE-open_chromatin model (5). Moreover, the online platform outcome only indicates the probability of the entire input sequence being a full, partial or non-open chromatin region without information on the chromatin status of specific regions within the input sequence. As a result, users will need to manually input and try overlapping subsequences multiple times to obtain more consistent and reliable predictions. Further, the prediction output consists of only text information (probability of being a full, partial or non-open chromatin region) without visualization of the DNA sequence region and its corresponding chromatin state.

To address this limitation, we developed the FveOCRs database, which adopts a sliding-window approach that at the same time utilizes the plant-dnamamba-BPE-open_chromatin model (5). The database automatically and systematically predicts the sequence’s chromatin state in a window when the window moves 10 bp at a time from 5’ to 3’. Further, predictions are for different genomic inputs, including the 5,000 bp upstream from the TSS, the first and second exons, the first and second introns, and the longest intron of the gene of interest. These regions were scanned with a small step size (10 bp) and varying window sizes (200bp, 500bp, and 1000bp) to estimate the probability of each window as a full, partial, or non-open chromatin region. The result from each step is shown as a dot in scatter plot or a line in JBrowse. This strategy delivers comprehensive and reliable results, allowing users to effectively identify potential open chromatin regions in the sequence of interest.

### Experimental validation

To experimentally evaluate the FveOCRs’ prediction, we tested the upstream sequence of the *FveMYB10* gene, encoding a transcription factor of anthocyanin pathway that promotes the red skin coloration of strawberry fruit (19). Analysis of the 5,000 bp upstream sequence (step size: 10 bp, window size: 1,000 bp) with FveOCRs predicted a full OCR located approximately 631-1700 bp upstream of *FveMYB10* (Figure 6C, D). Transient expression of a reporter gene *FveMYB10L* (20) was carried out in a white-fruited wild strawberry ‘Hawaii 4’ (21). The reporter was either driven by a 2.0 kb upstream sequence of *FveMYB10* containing a full OCR or by a 1.5 kb upstream sequence that has a truncated OCR (Figure 6E). The same 1.5 kb and 2.0 kb upstream sequences of *FveMYB10* were used to drive a GFP reporter as a non-pigment control. The *MYB10L* reporter driven by the full OCR (2.0 kb) gave a significantly higher level of reporter expression (in dark red) than the reporter driven by the truncated OCR (1.5 kb) (Figure 6E). The data supported the existence of a positive cis-element between 1.5 kb and 2.0 kb, which coincides with the presence of an OCR in the same region.

The *FveMYB10* example underscores the value of integrating the sliding-window approach with the plant-dnamamba-BPE-open_chromatin model. Within the 2,500 bp region upstream of *FveMYB10*, only a few windows (highlighted in brown) were predicted as full or partial open chromatin regions (Figure 6D). Therefore, sequence windows upstream or downstream of these brown dots have missed the detection of the open chromatin region upstream of *FveMYB10*. Therefore, entering sequences outside the brown dot region, if one were to use the PDLLMs online platform, would obtain non-open chromatin predictions. This limitation of the PDLLMs online platform is now overcome by the sliding window approach of FveOCRs.

In the future, we plan to further develop the FveOCRs database by offering more step-size and window-size options, incorporating additional genomic regions, and extending its scope to other Rosaceae species.

## Conclusions

We developed a database for predicting open chromatin regions in *Fragaria vesca* genome as well as in user-provided sequences of any plant species. The prediction is based on the sliding-window approach using the plant-dnamamba-BPE-open_chromatin model from PDLLMs. The database provides a systematic and precise OCR prediction in a useful visual output, which will facilitate cis-element identification for basic research and for engineering gene expression to improve crop traits.

## Data availability

The FveOCRs database is available at http://liulab.online:33838/FveOCRLiuLabTestVer25.

## Authors' contributions

M.L. and Z.L. conceived the research; M.L. and Z.Z. developed the database; Y.W. performed experiments; Z.L., S.M.M. and Q.M. supervised the research; M.L. and Z.L. wrote the manuscript.

## Acknowledgements

We would like to thank members of the Liu lab for providing helpful suggestions on the database.

